# Action potentials induce biomagnetic fields in Venus flytrap plants

**DOI:** 10.1101/2020.08.12.247924

**Authors:** Anne Fabricant, Geoffrey Z. Iwata, Sönke Scherzer, Lykourgos Bougas, Katharina Rolfs, Anna Jodko-Władzińska, Jens Voigt, Rainer Hedrich, Dmitry Budker

## Abstract

Upon stimulation, plants elicit electrical signals that can travel within a cellular network analogous to the animal nervous system. It is well-known that in the human brain, voltage changes in certain regions result from concerted electrical activity which, in the form of action potentials (APs), travels within nerve-cell arrays. Electrophysiological techniques like electroencephalography^1^, magnetoencephalography^2^, and magnetic resonance imaging^3,4^ are used to record this activity and to diagnose disorders. In the plant kingdom, two types of electrical signals are observed: all-or-nothing APs of similar amplitudes to those seen in humans and animals, and slow-wave potentials of smaller amplitudes. Sharp APs appear restricted to unique plant species like the “sensitive plant”, *Mimosa pudica*, and the carnivorous Venus flytrap, *Dionaea muscipula*^5,6^. Here we ask the question, is electrical activity in the Venus flytrap accompanied by distinct magnetic signals? Using atomic optically pumped magnetometers^7,8^, biomagnetism in AP-firing traps of the carnivorous plant was recorded. APs were induced by heat stimulation, and the thermal properties of ion channels underlying the AP were studied. The measured magnetic signals exhibit similar temporal behavior and shape to the fast de- and repolarization AP phases. Our findings pave the way to understanding the molecular basis of biomagnetism, which might be used to improve magnetometer-based noninvasive diagnostics of plant stress and disease.

---

Electrophysiological measurements enable investigation of the plant signaling pathways involved in reception and transduction of external stimuli, as well as communication within the plant body. Among the stimuli which can elicit pronounced plant electrical responses are light^9^, temperature^10^, touch^5,11^, wounding^12^, and chemicals^13^. In contrast to the three-dimensional complex electrical network of the human brain, the circuitry of a plant leaf is two-dimensional only. The bilobed trap of the *Dionaea* plant (Fig. 1a,b), formed by the modified upper part of the leaf, snaps closed within a fraction of a second when touched. Three trigger hairs that serve as mechanosensors are equally spaced on each lobe. When a prey insect touches a trigger hair, an AP (Fig. 1c) is generated and travels along both trap lobes. If a second touch-induced AP is fired within 30 s, the viscoelastic energy stored in the open trap is released and the capture organ closes^14^, imprisoning the animal food stock for digestion of a nutrient-rich meal. The leaf base, or petiole, is not excitable and is electrically insulated from the trap. Because of this, the trap can be isolated functionally intact from the plant by a cut through the petiole. On the isolated trap, mechanical stimuli trigger APs and closure just as on the intact plant. For the comfort of electrophysiological studies, one of the trap lobes can be fixed to a support while the other is removed, without affecting the features of AP firing. It has been shown that this simplified experimental flytrap system is well-suited to study the AP under highly reproducible conditions^15^. Other than by touch or wounding (mechanical energy), traps can be stimulated by salt loads (osmotic energy)^16^ and temperature changes (thermal energy).

**Fig. 1.**
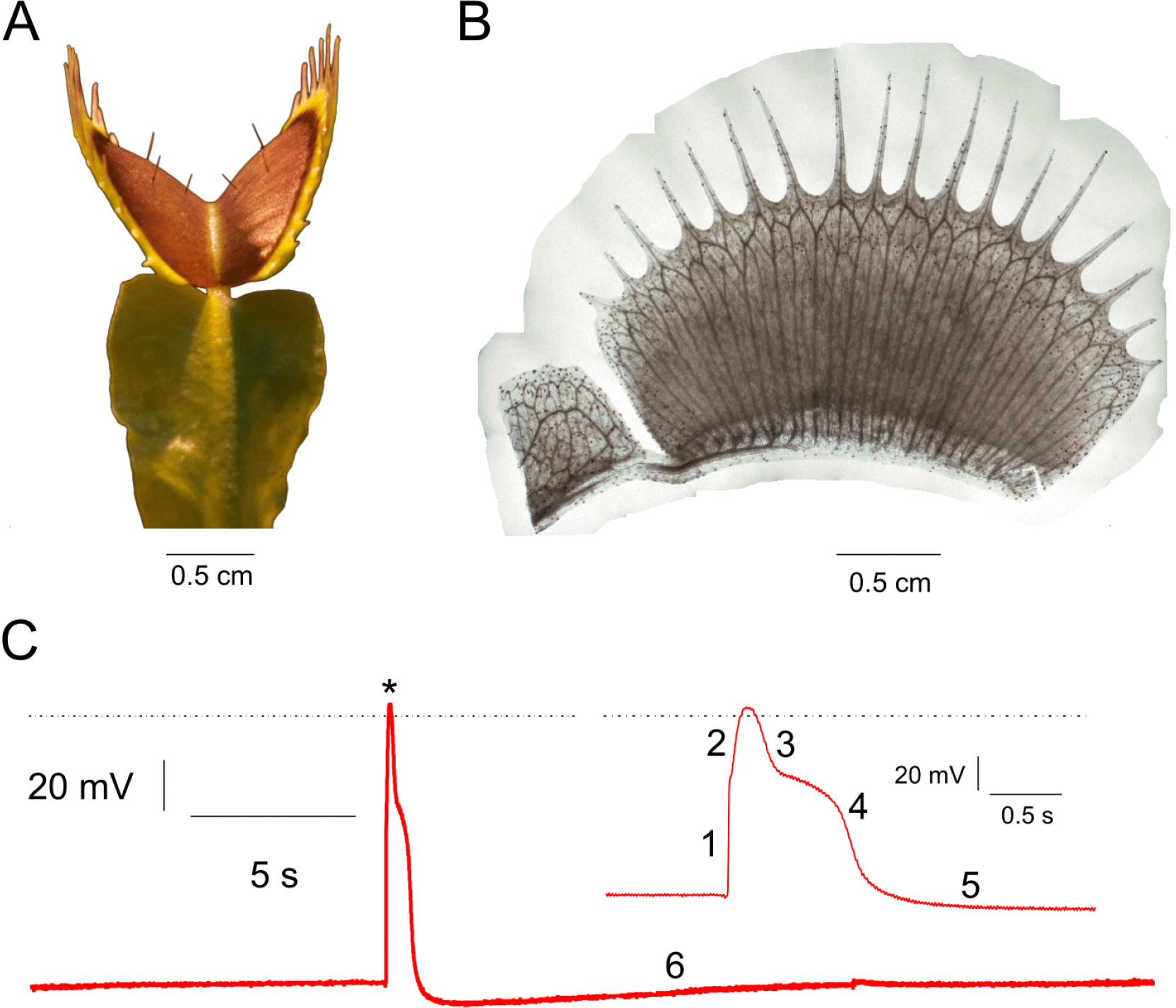
Venus flytrap geometry and action potentials. **a**, *Dionaea muscipula* leaf forms into a bivalved snap trap connected to the leaf base, or petiole. **b**, Side view of a single trap lobe showing vasculature structure. In contrast to the petiole, the trap contains parallel veins of interconnected cells. These veins consist of both dead low-conductivity water pipes (xylem) and living conductive phloem. Here the vasculature was imaged by staining for the dead vascular tissue. **c**, Intracellular AP lasting 2 s is subdivided into six phases (numbers), as explained in the text. The depolarization peak is indicated by an asterisk; the dotted line represents 0 mV. Inset, Zoom-in on the AP, resolving the first five phases of the AP.

Since touch activation of APs can cause unwanted mechanical noise in electric and magnetic recordings, we use thermal stimulation in our experiments. The interdisciplinary work presented here encompasses two complementary sets of experiments: the temperature dependence of flytrap electrical activity was studied in a plant-physiology laboratory, while magnetometer measurements of heat-stimulated traps were conducted in a magnetically shielded room.

## Heat-induced action potentials

When we heated up the support to which excised open traps were fixed, APs were elicited and the traps closed (Extended Data Fig. 1). To study the temperature dependence of heat-induced AP initiation, on one of the trap lobes we mounted a clamp equipped with a Peltier device and surface-voltage electrode (Fig. 2a). From a resting temperature of 20°C, the trap temperature was increased monotonically to 45°C at a rate of 4°C/s (Extended Data Fig. 2). Below 30°C, no APs were observed; above 30°C, the probability of AP firing increased and was maximal (100%) above 40°C. In 60 independent experiments using 10 different traps, we recorded the temperature at which an AP was first induced. When these data were plotted as temperature-dependent AP-firing probability (Fig. 2b), the curve could be well-fitted by a single Boltzmann equation characterized by a 50% AP-firing probability at 33.8°C. This behavior indicates that heat activation of the AP is based on a two-state process. The ion channels that carry the classical animal-type AP also occupy two major states: closed and open. In contrast to the animal sodium-based AP, the plant AP depolarization is operated by a calcium-activated anion channel^5^. Thus, we conclude that the temperature “switch” of the *Dionaea* AP is based on a calcium-dependent process. Following Ca^2+^ binding, the anion-channel gates open. Our experiments indicate that at temperatures of *T* ≲ 34°C the cellular Ca^2+^ level remains below threshold, but at T ≳ 34°C there is enough chemical energy to open a critical number of anion channels, driving the fast depolarization phase of the AP.

**Fig. 2.**
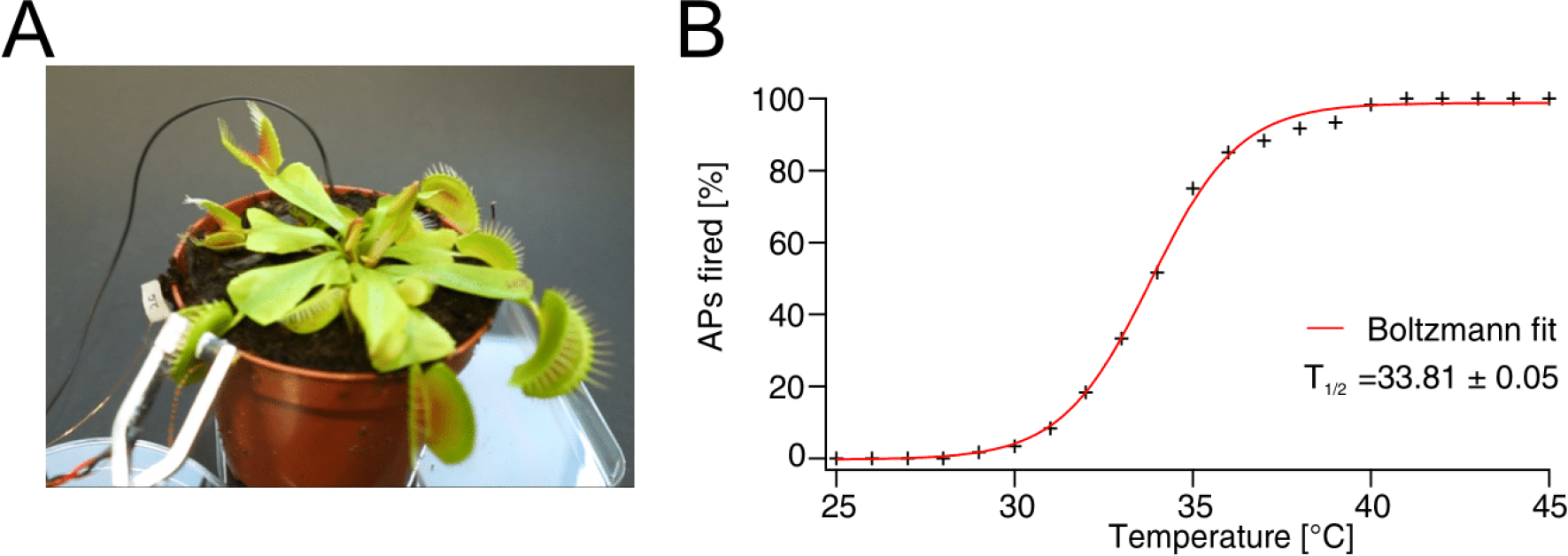
Electrical measurements of heat-induced action potentials. **a**, *Dionaea* plant with clamp mounted on one lobe of a trap, equipped with a Peltier device and surface-voltage electrode. A ground electrode is placed in the soil surrounding the plant root. **b**, Temperature dependence of AP-firing probability fitted by a Boltzmann equation (red curve), characterized by 50% firing probability at temperature *T*_1/2_.

The *Dionaea* AP can be subdivided into 6 well-defined phases (Fig. 1c): (1) fast depolarization, (2) slow depolarization, (3) fast repolarization, (4) slow repolarization, (5) transient hyperpolarization, and (6) slow recovery of the membrane potential to the pre-AP state. When comparing APs recorded at different temperatures, we found that temperature affects the signal amplitude and duration. Increasing the thermal energy input changed not only the probability for an AP to be fired, but also led to an increased AP amplitude and decreased half-depolarization time (Supplementary Information). These facts indicate that heat-sensitive ion channels trigger and shape the AP: at higher temperatures, thermal energy input causes more closed Ca^2+^-activated anion channels to open and depolarize the membrane potential. Compared to depolarization, fast repolarization (mediated by K^+^ channels) and transient hyperpolarization (caused by depolarization activation of outward-directed protein pumps) were much less affected by temperature. The recovery time to reach the resting membrane potential was essentially insensitive to temperature changes.

Besides lowering the AP firing threshold and changing certain features of the AP, prolonged heat stimulation can induce trap lobes to enter an autonomous AP firing mode (Extended Data Fig. 1). When increasing the bottom surface temperature of the recording-chamber base from 20 to 46°C, AP spiking activity sets in after a couple of seconds, reaching a steady AP firing frequency of 3.8 per minute at a stable 46°C surface temperature. Induction of autonomous APs has also been obtained using flytraps treated with NaCl salt (osmotic energy)^17^.

## Biomagnetism

Having established heat stimulation as a reliable noninvasive technique for inducing flytrap APs, we searched for the magnetic field associated with this electrical excitability. Magnetometry experiments were carried out at Physikalisch-Technische Bundesanstalt (PTB) Berlin in the Berlin Magnetically Shielded Room 2 (BMSR-2) facility^18^, using four QuSpin Zero-Field Magnetometers (QZFM). These commercial optically pumped magnetometers (OPMs) employ a glass cell containing alkali vapor to sense changes in the local magnetic-field environment^8^. A magnetically shielded environment is required for operation of the magnetometers, and use of a walk-in shielded room allowed for the constant presence of an experimenter to prepare plant samples and carry out measurements. As shown in Fig. 3, an isolated trap lobe was attached to the housing of the primary sensor (denoted A), such that the distance between the plant sample and the center of the atomic sensing volume was approximately 7 mm. Two secondary sensors (B and C) were placed nearby the primary sensor to measure signal fall-off, and an additional background sensor (D) was used to monitor the magnetic environment in the shielded room. Each magnetometer is sensitive to signals along two orthogonal axes. Sensor electronics were connected to a data-acquisition system in the PTB control room outside the magnetically shielded room. To monitor heat-induced APs, we used two silver-tipped copper surface electrodes, inserted in either end of the plant sample^19^. These data, together with other auxiliary trigger signals, were sampled simultaneously with the OPMs using the same data-acquisition system.

**Fig. 3.**
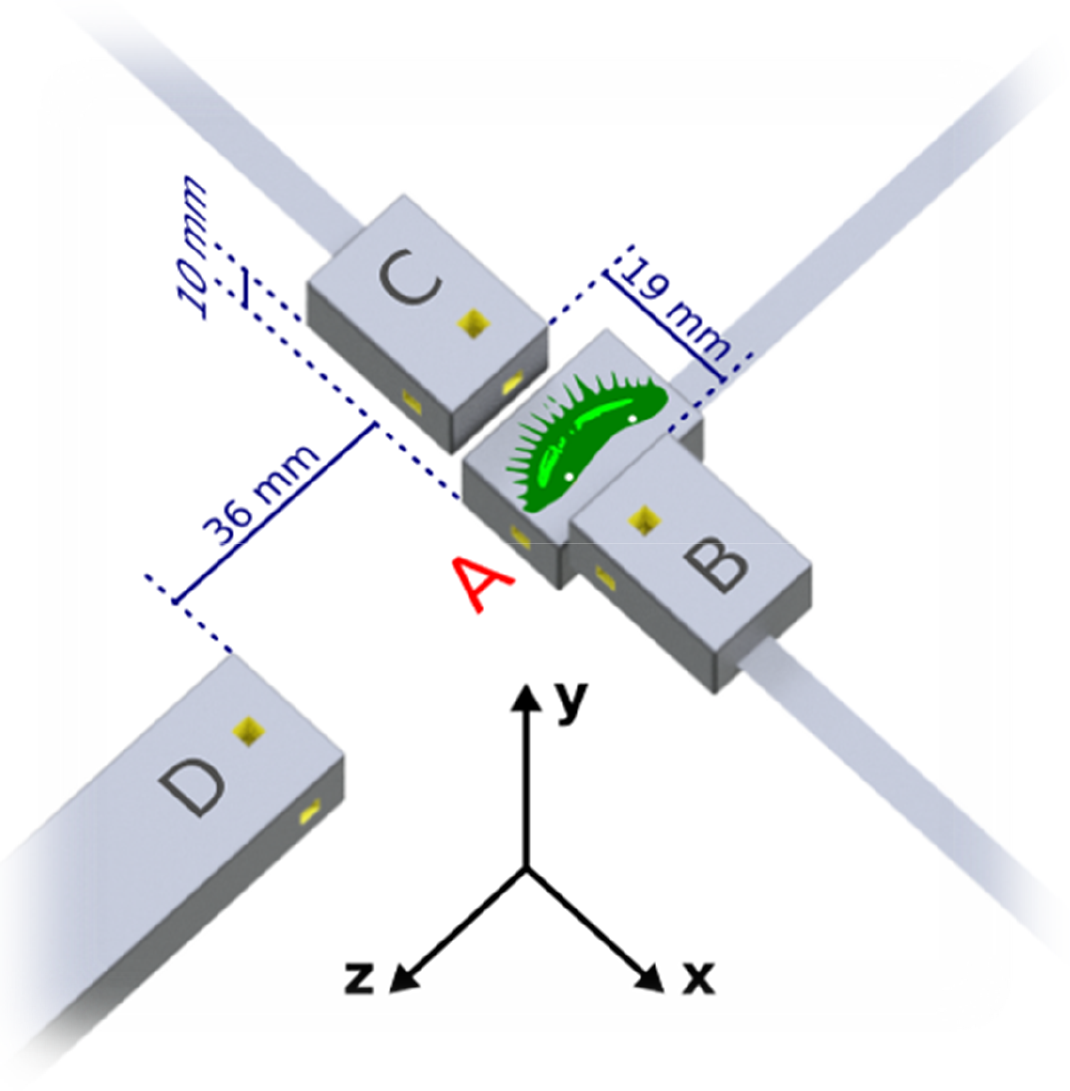
Schematic of the experimental setup in the magnetically shielded room. The plant sample, an isolated lobe of the flytrap, is placed on top of primary sensor A, in the *x-z* plane with trigger hairs exposed. For reference, the dimensions of the housing (gray boxes) for the primary and secondary sensors are 24.4×16.6×12.4 mm^3^. Yellow cut-outs indicate the position of the 3×3×3 mm^3^ atomic sensing volume. A 3D-printed ABS plastic structure (not shown) holds the magnetometers in position on a wooden table. White dots on the plant sample, approximately 1 cm apart, indicate the placement of the surface electrodes for AP monitoring. In the coordinate system shown, all magnetometers are sensitive along the *y*-axis, normal to the surface of the plant sample; furthermore, A and D are sensitive along the *z*-axis, and B and C are sensitive along the *x*-axis. Sensors B and C are positioned symmetrically around sensor A. Sensor D serves as a background sensor and is therefore located farther away from the sample.

Resistive heaters in the magnetometer housing, which are used to increase the atomic density and improve sensitivity, also served to induce autonomous AP firing via surface heat transfer. Electric and magnetic signals were recorded simultaneously from traps heated to a surface temperature of 41°C. Prior to the measurements, we performed tests to ensure that no spurious magnetic fields were generated by the electrode system (Supplementary Information). To better distinguish the observed magnetic signals from background noise, we triggered on the electric signals and averaged the magnetic data in a time window around those trigger points. Examples of averaged magnetic data are shown in Fig. 4a,b. A clear magnetic signal with a time scale corresponding to that of the averaged electric signal is visible in the primary-sensor data. For comparison, data from several different experiments were plotted (Fig. 5). To minimize common background noise, we subtracted the magnetic data of sensor D to create a gradiometer with a 48-mm baseline. Signals of up to 0.5 pT are visible in the *y*-axis gradiometric data, normal to the sample surface. The signal magnitude obtained was comparable to what one observes in surface measurements of nerve impulses in animals^20^.

**Fig. 4.**
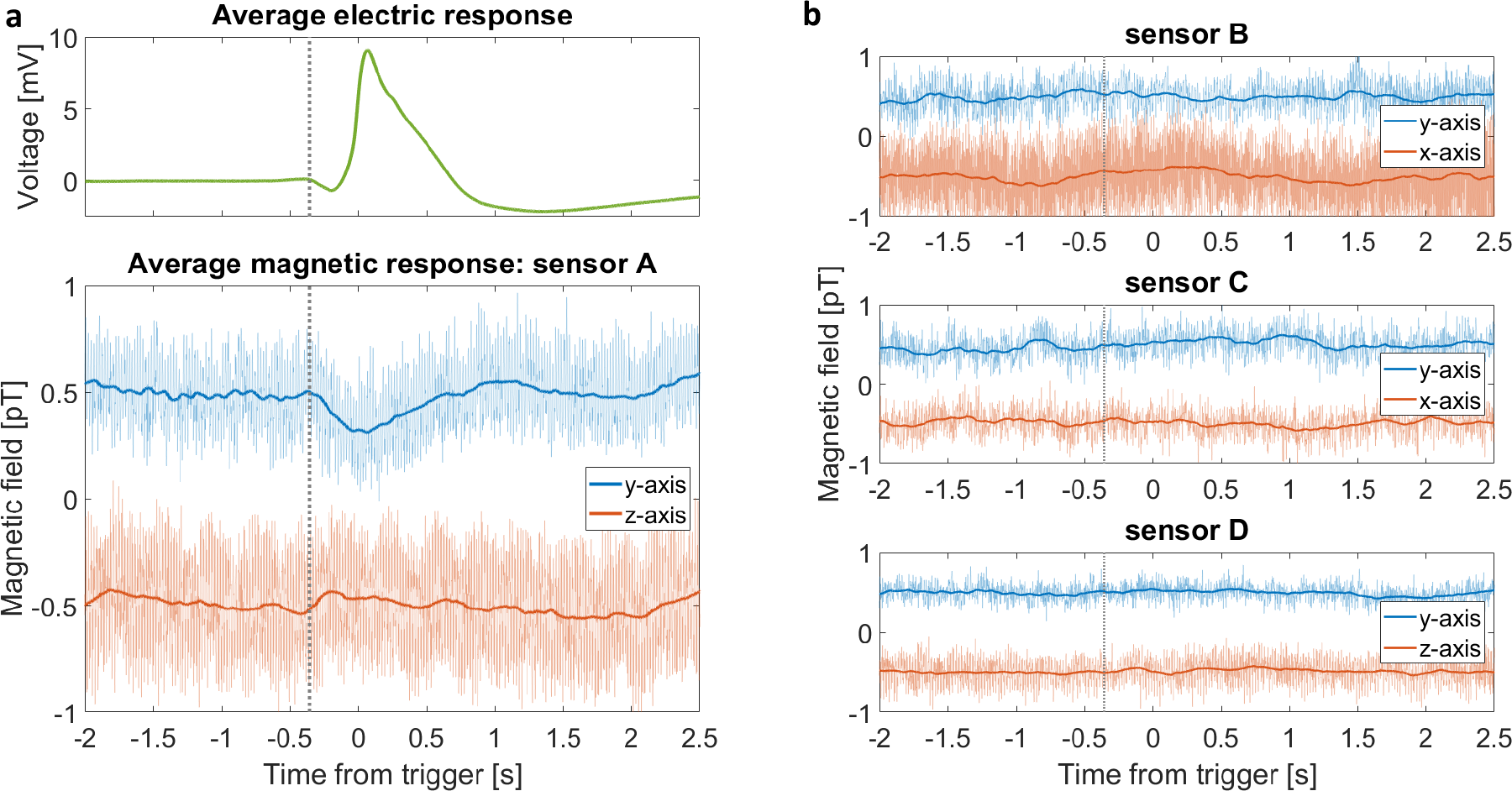
Average action potential and corresponding magnetic signals. **a**, Result of triggering on nine consecutive APs from a trap lobe heated to 41°C, then averaging the electric and magnetic data from a 4.5 s window around each trigger point. The average magnetic traces (bottom graph, opaque traces) were frequency-filtered (50 Hz low-pass), then smoothed with a 0.2 s running average. A magnetic signal is visible in both sensitive axes of the primary sensor A. For comparison, the raw unfiltered data are plotted behind the processed data. For visual clarity, DC offsets have been added to the data, and vertical gray dotted lines indicate the approximate start time of the electric signal. **b**, Average magnetic response from the other three sensors, obtained using the same procedure as in **a**. The data from the secondary sensors, B and C, do not show a signal. The data from the background sensor D can be used to remove noise common to all sensors (see Fig. 4).

**Fig. 5.**
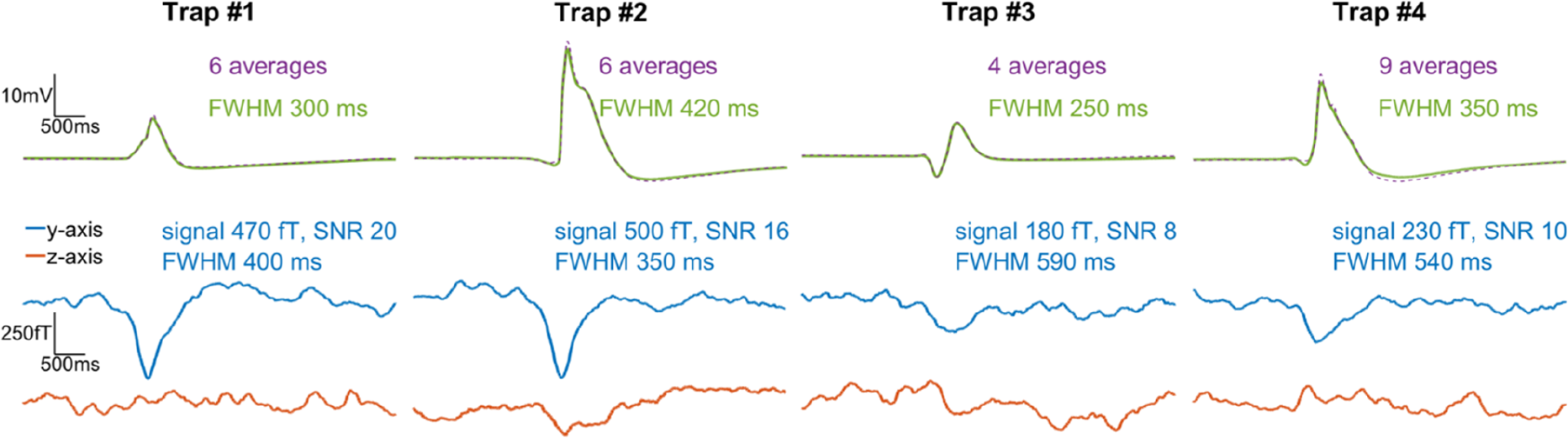
Comparison of average electric and gradiometric signals from four different experiments. In each case we triggered on heat-induced APs in the electric trace and performed the same data analysis as for Fig. 4. The electric response recorded by the surface electrodes (top row; average signal plotted as solid green, single AP plotted as dashed purple) varies in amplitude because a different plant sample was used in each experiment. To produce the gradiometric plots (bottom graphs) we subtracted the magnetic data of background sensor D from that of primary sensor A. The number of averages in each experiment is indicated, along with the amplitude and signal-to-noise ratio (SNR) of the *y*-axis gradiometric signal. The rightmost panel (Trap #4) shows the same data set as in Fig. 4.

To quantify the significance of the signals, signal-to-noise ratios (SNRs) were calculated from the average *y*-axis gradiometric time traces in Fig. 5 as follows. The noise level is defined as the standard deviation of the gradiometric response in a 1.5 s time window (from time *t* = −2 s to *t* = −0.5 s in Fig. 4a) prior to signal onset. The signal size is defined as the amplitude of the extreme (minimum) field value, with respect to the mean value in the noise window. For the four experiments shown in Fig. 5, the SNRs range from 8 to 20. The corresponding p-values are *p* < 9 × 10^−16^, indicating that the probability of such signals arising from random noise is negligible. At the sub-Hz signal frequency, the sensitivity of the gradiometer is approximately 100 fT/√Hz (Extended Data Fig. 6). For both the electric and magnetic signals, the full width at half extremum (maximum or minimum, FWHM) were also calculated, where the extremum is defined with respect to the mean value in the noise window.

The temporal superposition of the electric and magnetic signals in Fig. 5 suggests that we have detected the magnetic activity associated with the flytrap AP. Unlike in measurements of animal nerve axons and the large internodal cells of *Chara corallina* alga, where the magnetic field is proportional to the time derivative of the intracellular voltage^21,22^, the magnetic signal from the complex multicellular flytrap lobe has a shape similar to the electric signal. We see features in the magnetic signal which appear to correspond to the depolarization and repolarization phases of the AP. In electric recordings using surface electrodes, the exact shape and duration of signals are dependent on the placement of electrodes on the measured sample. By contrast, magnetometry records a “true” physical signal from the organism. In this sense, it is comparable to intracellular electrode techniques. Whereas intracellular electrodes are sensitive to electrical activity of single cells, magnetometers can record both local and systemic activity at the multicellular level.

The physical origin of the measured biomagnetic fields is related to an outstanding question in plant electrophysiology: how electrical signals propagate over long distances through the plant. Essentially this is a scaling problem: while electrical signaling is well-understood in some unicellular plant systems^22^, much less is known about the propagation mechanisms of such signals between cells and along cellular pathways. For the Venus flytrap system, it is known from electrode measurements that APs propagate through the trap at speeds of around 10 m/s ^6^. A proposed pathway of long-distance signal propagation between plant cells in the trap is the electrically conductive phloem in the vasculature (Fig. 1b). Given that the typical resistance between two points on a trap is *R* ≈ 1 MΩ ^23^, we can perform a basic calculation to confirm that the magnitude of the magnetic fields we measure is reasonable. We estimate the expected magnetic-field magnitude at the center of the sensing volume to be

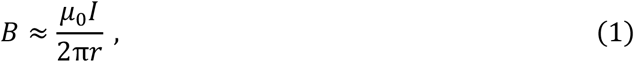

where *I* = *V/R* ≈ 10 nA is the current passing through the trap between the electrodes, and *r* ≈ 7 mm is the perpendicular distance from the trap surface. Using these values, we find *B* ≈ 0.3 pT, a magnitude which corresponds well with the *y*-axis experimental results of sensor A. Although the precise distribution and directionality of current flow in the trap is unknown, we can use the geometry of the trap (Fig. 1a,b) and magnetometry setup (Fig. 3) to further interpret our results. If the *x*-oriented parallel-cable structure of the vasculature is the primary conduction pathway, magnetic field along the *y*-direction is expected at the primary sensor A, but not at the secondary sensors B and C. The symmetry of the trap about the *x*-direction could explain the relative lack of *z*-axis magnetic signal in our measurements. (Trap curvature and misalignment with respect to the sensor housing may give rise to *z*-axis signals in some experiments.) Thus, our magnetometry results agree with a hypothesis that the vasculature serves as a network for long-distance electromagnetic signaling within the trap.

## Discussion

Although human and animal biomagnetism are well-developed areas of research^2,20,21,24,25,26^, very little analogous work has been conducted in the plant kingdom^9,12,22,27^, largely because biomagnetic signals are typically much smaller in amplitude and frequency than their animal counterparts. Previously reported detection of plant biomagnetism, which established the existence of measurable magnetic activity in the plant kingdom, was carried out using superconducting-quantum-interference-device (SQUID) magnetometers^9,12,22^. Atomic magnetometers are arguably more attractive for biological applications, since, unlike SQUIDs^28,29^, they are non-cryogenic and can be miniaturized to optimize spatial resolution of measured biological features^20,26,30^. Our study of plant biomagnetism using atomic magnetometers documents: (i) existence and features of biomagnetic signals in the Venus flytrap, (ii) magnetic detection of APs in a multicellular plant system generally, and (iii) electric and magnetic detection of heat-induced APs in the Venus flytrap. In the future, the SNR of magnetic measurements in plants will benefit from optimizing the low-frequency stability and sensitivity of atomic magnetometers. Just as noninvasive magnetic techniques have become essential tools for medical diagnostics of the human brain and body, this noninvasive technique could also be useful in the future for crop-plant diagnostics—by measuring the electromagnetic response of plants facing such challenges as sudden temperature change, herbivore attack, and chemical exposure.

