## Supplementary material for "Action potentials induce biomagnetic fields in Venus flytrap plants": Methods, Supplementary information, Extended data figures

To obtain strong electric and magnetic signals, the health of the plants is paramount. We purchase adult Venus flytraps from a carnivorous-plant greenhouse (Gartenbau Weilbrenner, Freinsheim, Germany). Normally the plant samples are housed in a growth chamber manufactured by Poly Klima. To keep the flytraps alive during the PTB measurement run, we used homemade plastic greenhouses equipped with plant-cultivation lighting and temperature and humidity monitoring. The plants were kept on an automated 12/12-hour light/dark cycle at approximately 25°C and 75% relative humidity, treated only with distilled water.

For recording of flytrap APs in our heat-stimulation investigations, we used surface electrodes measuring the extracellular potential of a trap. One silver electrode was inserted into the trap, with the electrical connection enhanced by application of a droplet of contact gel (Laboklinika), while the reference electrode was inserted into wet soil or the petiole midrib. Electrical signals were amplified 100-fold and recorded with Patchmaster software (HEKA). Temperature dependence of AP induction was studied by application of a homemade Peltier device powered by a PTC-10 temperature-control system (npi electronic, NJ 08510, United States). Constant heat was applied using an IKA RET basic hot plate (IKA-Werke GmbH & Co. KG, Staufen, Germany) heated to 46°C.

Several types of magnetometry experiments were conducted at PTB: controls, OPM measurements using four QuSpin sensors (three Gen-2: denoted A, B, C; one Gen-1.5: denoted D), and measurements using the multi-channel SQUID array of BMSR-2. See Supplementary Information for further details of the SQUID measurements.

For the OPM measurements of isolated trap lobes, each sample was cleaved from the plant with a razor blade and placed on the primary sensor A for immediate measurement. The sample was either secured to the sensor housing with double-sided adhesive tape (acrylate, thickness 0.5 mm) or placed on a plastic slide (PET, thickness 0.22 mm) on the housing. Electrode, magnetometer, and electric reference signals were recorded at a 500-Hz acquisition rate on a 9-channel analog data-acquisition system with PC control. The raw difference signal from the two surface electrodes was first sent through a voltage preamplifier (Stanford Research Systems, Model SR560), AC-coupled with a gain of 100. It is essential to use a voltage, rather than current, preamplifier to avoid currents in the electrical leads whose magnetic fields may be detected by the magnetometers. Since leakage of electrical signals into magnetic channels is a serious concern, we address the topic in detail in Supplementary Information.

### Data availability

The datasets generated and analyzed during the current study are available from the corresponding author on reasonable request.


### Acknowledgements

We acknowledge the support of the Core Facility "Metrology of Ultra-Low Magnetic Fields" at Physikalisch-Technische Bundesanstalt, which receives funding from the Deutsche Forschungsgemeinschaft (DFG KO 5321/3-1 and TR 408/11-1). Dr. Tilmann Sander-Thömmes and Sophia Haude assisted during the PTB data run. Dr. Rob Roelfsema of the University of Würzburg provided valuable guidance in the early stages of the project. Pavel Fadeev offered helpful comments on the manuscript. A.F. was supported by a Carl-Zeiss-Stiftung graduate fellowship. This research was supported in part by the German Federal Ministry of Education and Research (BMBF) within the Quantumtechnologien program (FKZ 13N14439), as well as the DFG Koselleck award HE 1640/42-1 to R.H.


### Author contributions

A.F. and D.B. proposed to study biomagnetism in the Venus flytrap. A.F., G.I., S.S., L.B., K.R., A.J.W., and J.V. conducted experiments. A.F. and S.S. analyzed data. A.F., R.H., and S.S. wrote the manuscript. R.H., D.B., G.I., L.B., S.S., J.V., A.J.W., and K.R. edited the manuscript. D.B. and R.H. supervised research.

### Competing interest declaration

The authors declare no competing interests.

### Supplementary information

As part of our study of heat-induced flytrap electrical behavior, we compared the amplitude and depolarization kinetics of APs recorded at 10, 20, 30, and 40°C. There was a 1.6-fold increase in AP amplitude from 10 to 40°C. When heating the trap from 10 to 30°C, the half-depolarization time dropped from $0.29 \pm 0.08$ s to $0.13 \pm 0.02$ s.

In the PTB data run, as a complement to the OPM measurements we also conducted two types of experiments using 57 channels of the BMSR-2 built-in SQUID array. The first type involved placing an intact flytrap plant directly under the SQUID dewar, whose bottom surface has a 2.8-cm offset from the plane of the pick-up coils. We closed each trap in turn by two consecutive mechanical stimulations of the trigger hairs with a plastic pipette tip. In the data analysis, we looked for signals in the magnetic data corresponding to either the APs or subsequent trap closure. Even after averaging multiple SQUID channels, no signals were found, probably because of the large distance between sample and sensors. In the second type of SQUID experiment, we attached an isolated trap lobe directly to the bottom of the dewar and performed mechanical stimulation, but again no magnetic signals were found during data analysis. Following the calculation in the main body of the paper, at offset distance of at least 2.8 cm from the sample we would reasonably expect a magnetic-field magnitude on the order of 10 fT. Since this is approximately the noise floor of the SQUID magnetometer system at 1 Hz (under ideal operating conditions), the null result is consistent with expectations.

Prior to the data run, we tested the electrode system to ensure that no spurious magnetic fields due to currents in the electrode wires would be picked up by the magnetometers under usual experimental conditions. These tests were conducted using the circuit depicted in Extended Data Fig. 3, with a four-layer MS-2 magnetic shield from Twinleaf containing two active QuSpin sensors (B and C) placed side-by-side. A function generator (Tektronix AFG2021) in parallel with a resistor created a sawtooth "artificial flytrap action potential" signal at 1.2 Hz. This signal was sent through a low-noise voltage preamplifier (SRS Model SR560) with typical experimental settings (6-dB low-pass 10-Hz filter, AC coupling, gain 1000, input impedance 100 MΩ) that yielded a preamplifier output of amplitude 2 V—corresponding to the output amplitude we would see in an actual flytrap experiment. The crucial step was to simulate a "worst-case scenario" for the electrode wires. To that end, a 1-cm copper coil, in series with the preamplifier, was placed directly on top of sensor C. As is evident in Extended Data Fig. 4, no signal at 1.2 Hz was visible in the $y$-axis or $z$-axis data of either magnetometer. Even when we increased the amplitude of the electric signal by five times (corresponding to 10-V preamplifier output), no signal at 1.2 Hz was observed. Thus, we were satisfied that the electrode/voltage-preamplifier system was not a source of unwanted noise. The voltage preamplifier and electronics used in these diagnostic experiments were the same as those used in BMSR-2 for plant experiments.

For comparison, we also conducted identical tests with a low-noise *current* preamplifier (SRS Model SR570, 6-dB low-pass 10-Hz filter, sensitivity 100 nA/V). In this case the signal at 1.2 Hz did appear above the noise in the data of sensor C. For example, the 2-V experiment yielded a 3-pT signal along the $y$-axis, indicating that a current of over 20 nA was flowing in the current loop. Based on these results, we exclusively used the voltage preamplifier in our data run at PTB. As an additional security check, in all OPM experiments we ran one of the electrode wires over the background sensor D to monitor for possible spurious signals (none were detected).

Extended Data Fig. 5 shows the electric time traces used in the data analysis for Fig. 3 and Fig. 4 in the main text. The recorded APs are slightly variable in shape and exhibit certain artifacts, which is normal for surface-electrode measurements. In some time traces (e.g. Extended Data Fig. 5a) we observed the frequency of autonomous AP firing increasing over time, which may be explained as follows. Sufficient input energy is required to increase the cytosolic calcium level to threshold—once this threshold is reached, an AP is released. As the trap heats up in our setup, the stored cellular energy increases while the new energy which needs to be input for the next AP decreases, which could lead to an increase in AP firing frequency.

To characterize the performance of the QuSpin gradiometer system in the shielded room, we recorded the background in the room and performed frequency analysis. A typical noise spectrum is shown in Extended Data Fig. 6.


**Corresponding author**

Correspondence to Anne Fabricant.

### Extended data figures

a

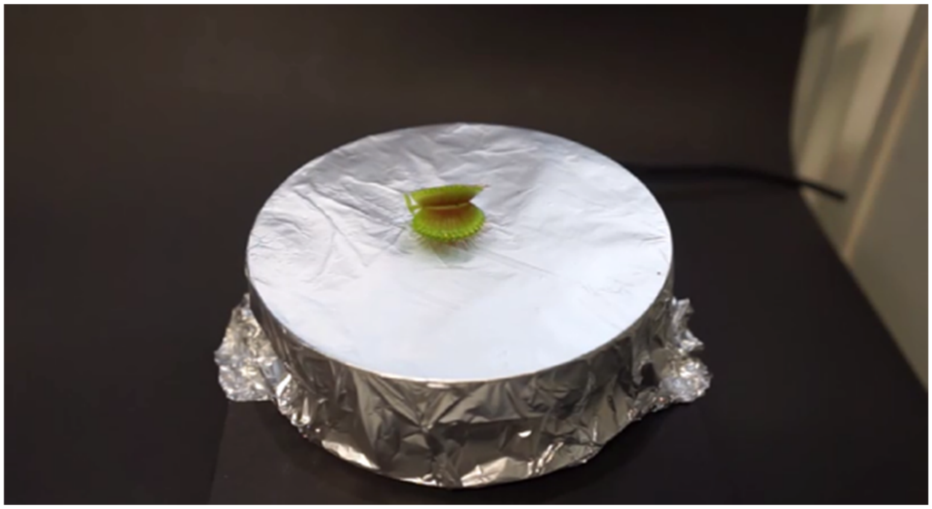

b

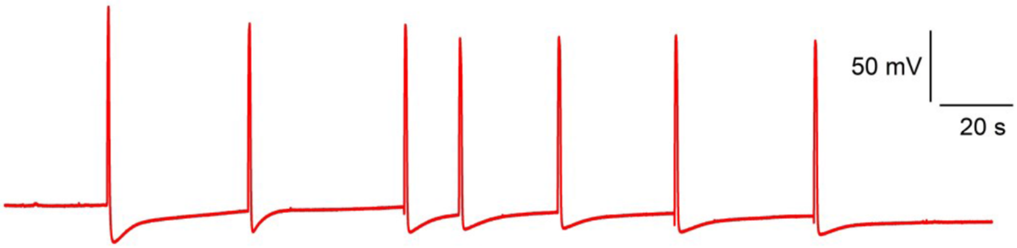


**Extended Data Fig. 1 | Spontaneous AP firing on a hot plate heated to 46°C. a**, Video (online) showing trap closure on the hot plate; playback speed is increased by a factor of 10. **b**, Surface-potential measurements of the heated trap, confirming that heat evokes APs.

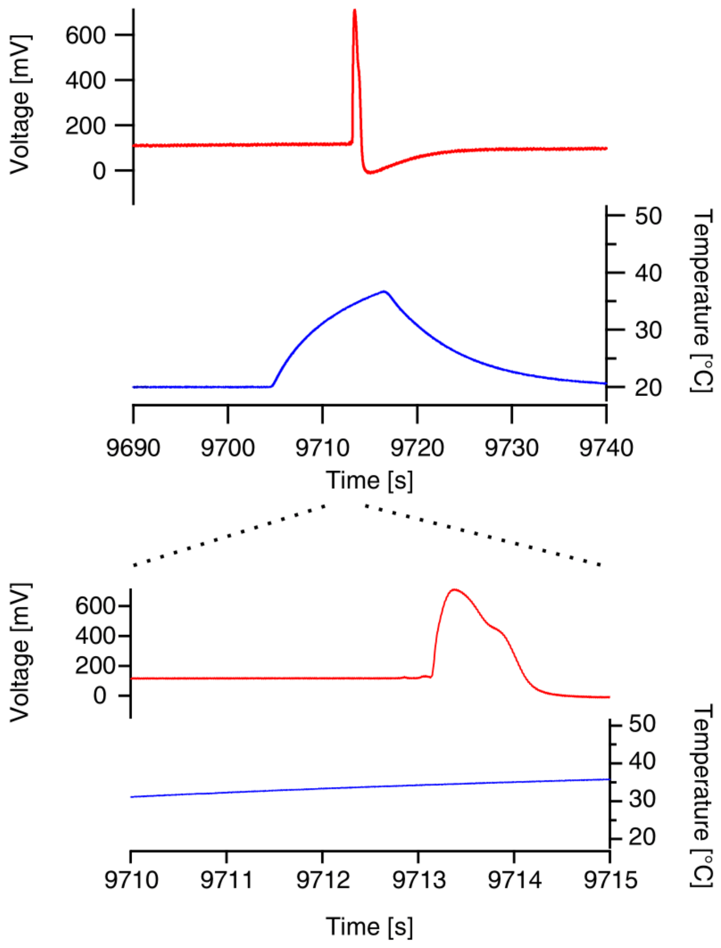


**Extended Data Fig. 2 | Comparison of measured surface potential and applied temperature.** Heat was applied via a Peltier element placed on the inner trap surface. An AP (red curve) occurred as the temperature (blue curve) increased from 20 to 45°C. The lower graph is a zoom-in on the time axis to define the temperature at which the AP occurred.

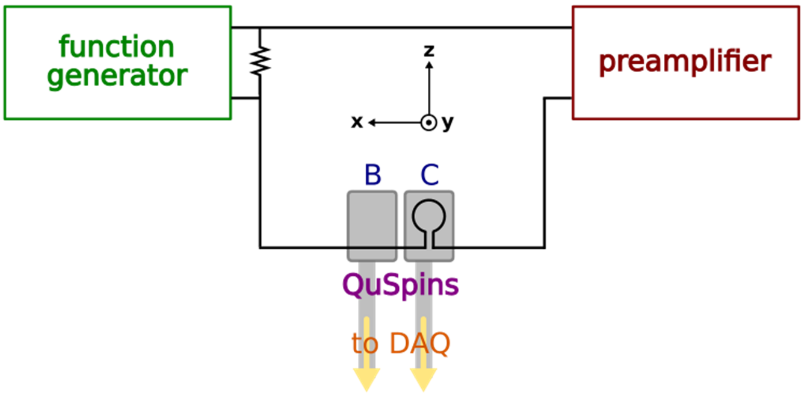


**Extended Data Fig. 3 | Circuit for testing the preamplifiers.** See Supplementary Information for details.

**a**

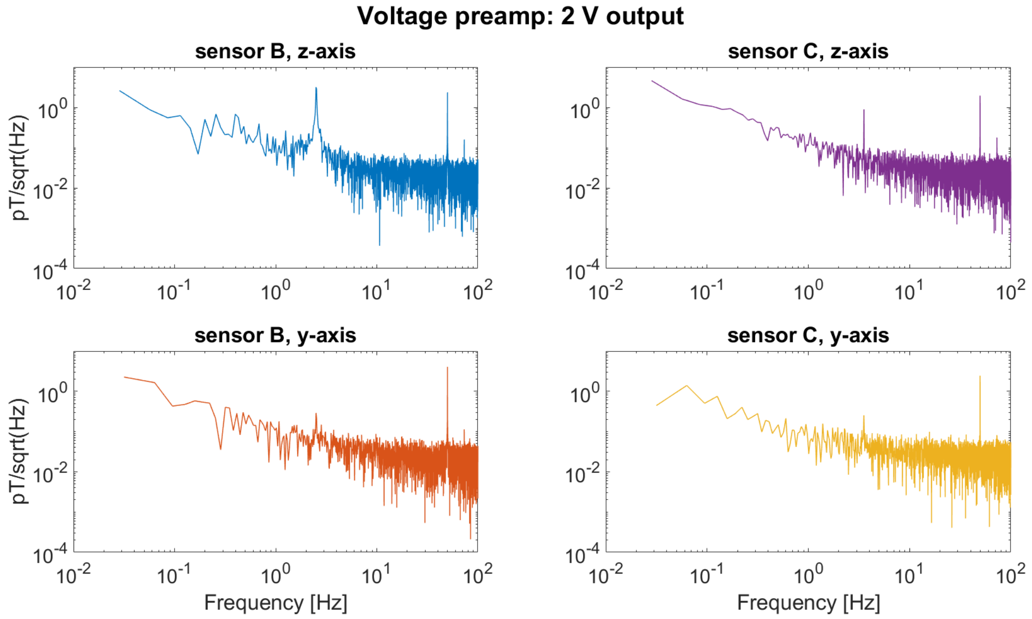
Voltage preamp: 2 V output
sensor B, z-axis
sensor C, z-axis
sensor B, y-axis
sensor C, y-axis
pT/sqrt(Hz)
Frequency [Hz]

**b**

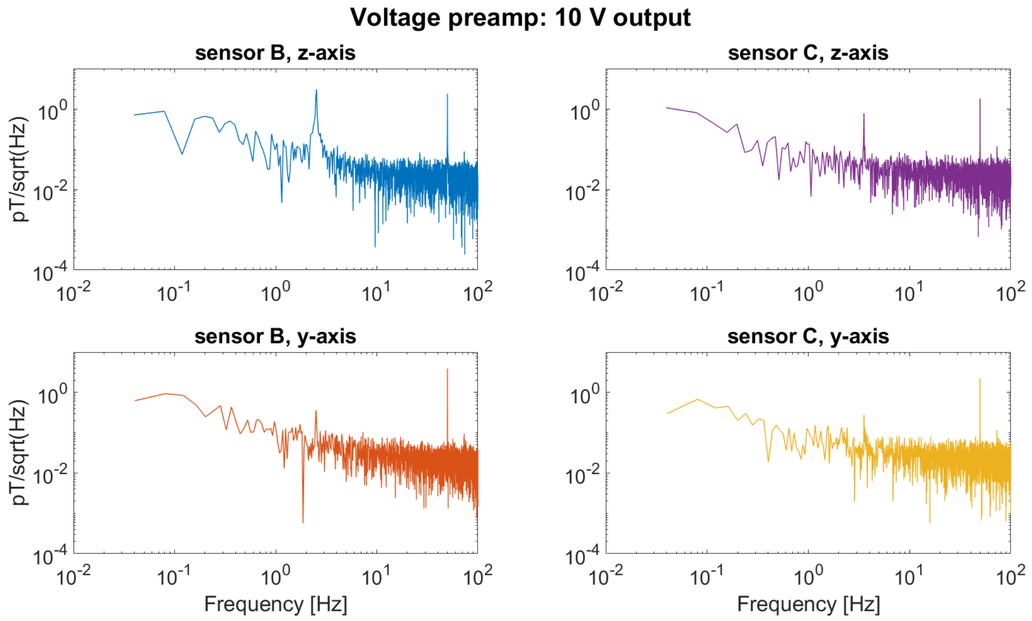
Voltage preamp: 10 V output
sensor B, z-axis
sensor C, z-axis
sensor B, y-axis
sensor C, y-axis
pT/sqrt(Hz)
Frequency [Hz]

**c** **d**

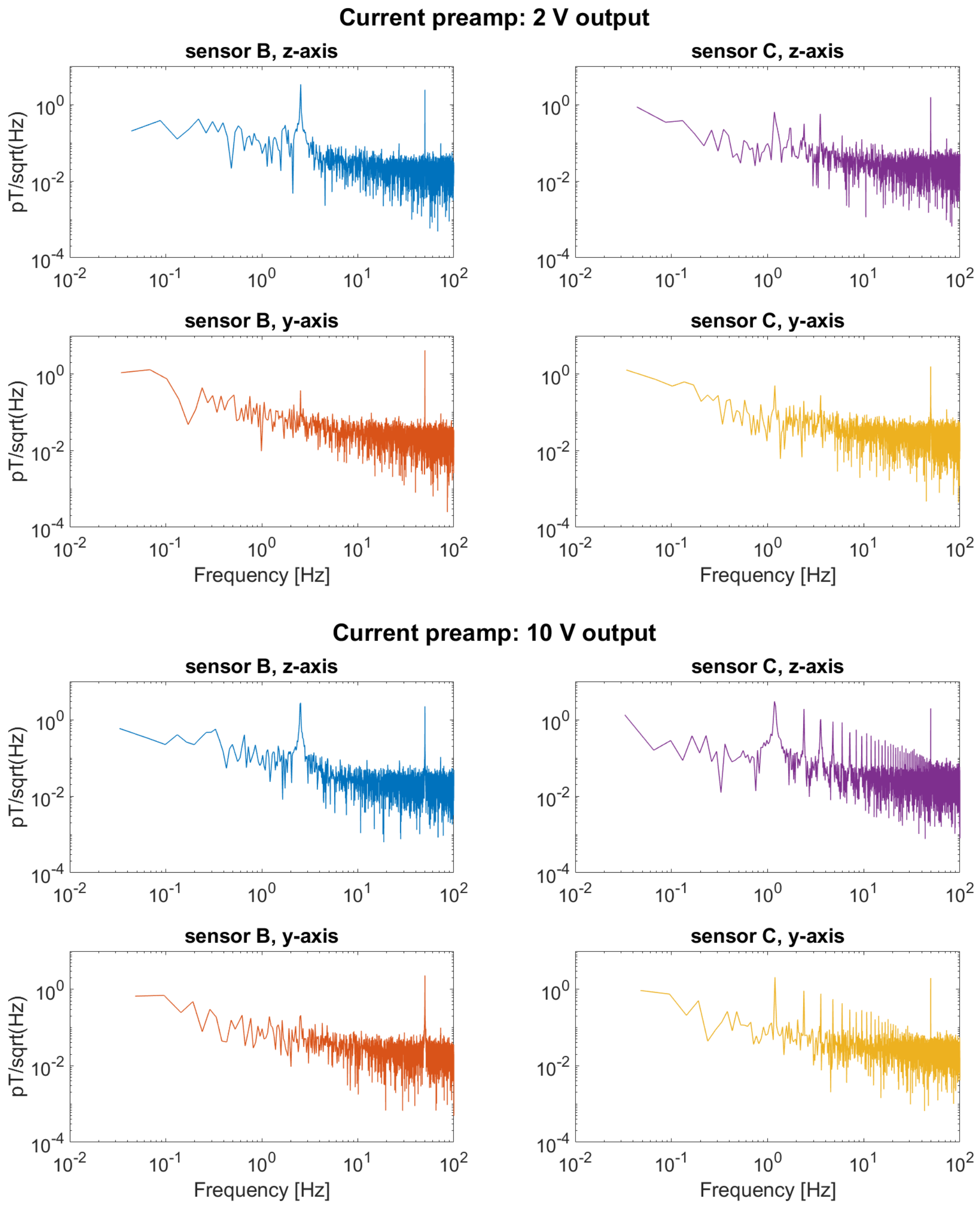


**Fig. S4 | Results of the preamplifier tests.** In addition to the 50-Hz line frequency, peaks due to lab background noise appear at 2.5 and 3.5 Hz. Signal from the current preamplifier appears in the data of sensor C (**c** and **d**), but this effect is not seen when the voltage preamplifier is used (**a** and **b**).

a

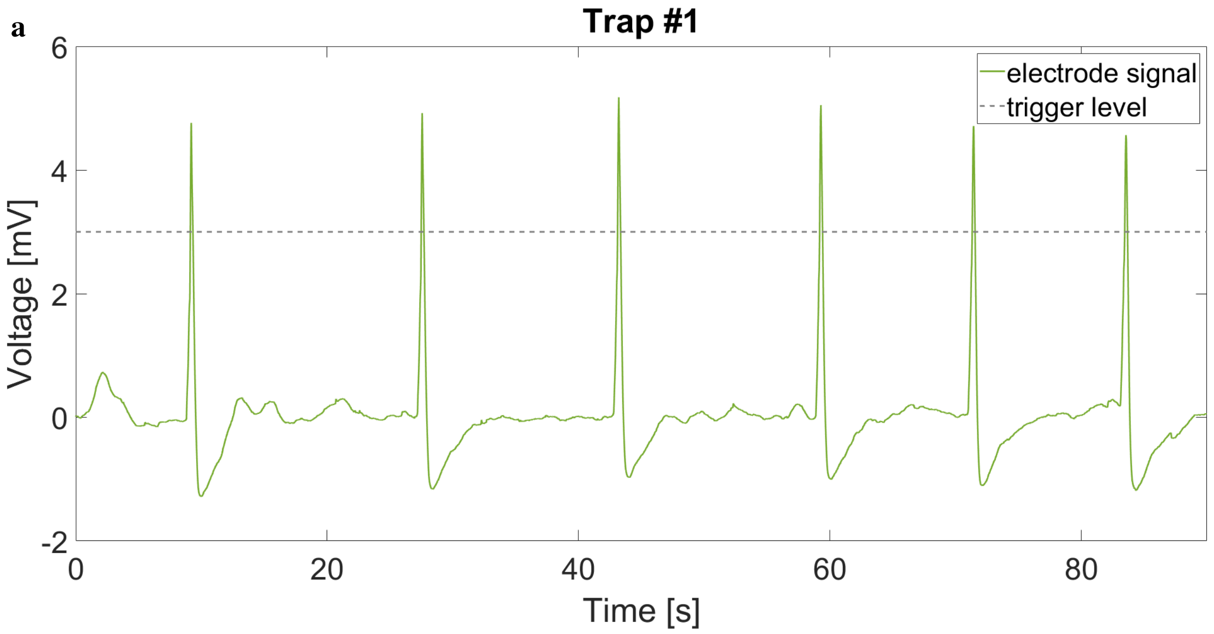

Trap #1
electrode signal
trigger level
Voltage [mV]
Time [s]
6
4
2
0
-2
0
20
40
60
80


b

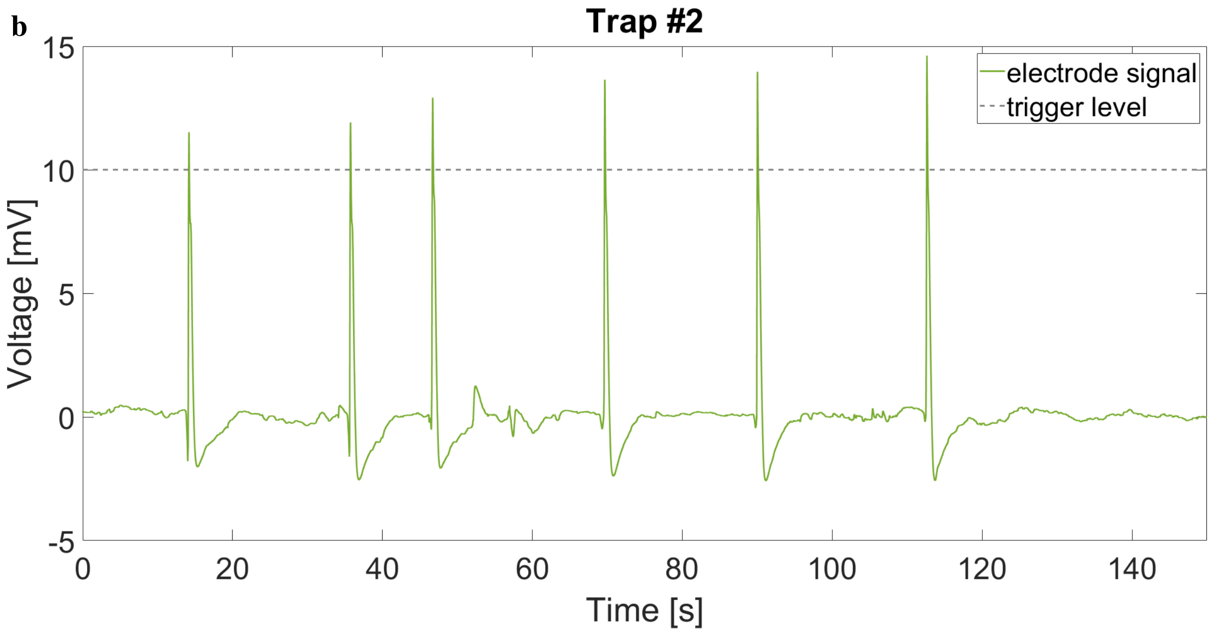

Trap #2
electrode signal
trigger level
Voltage [mV]
Time [s]
15
10
5
0
-5
0
20
40
60
80
100
120
140

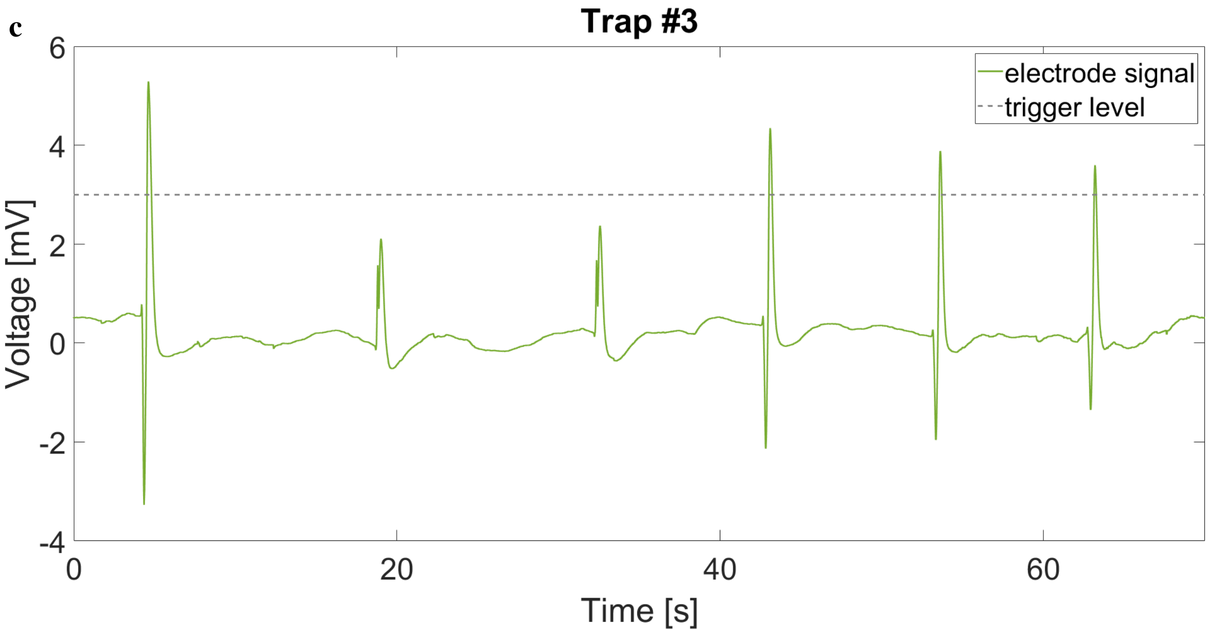


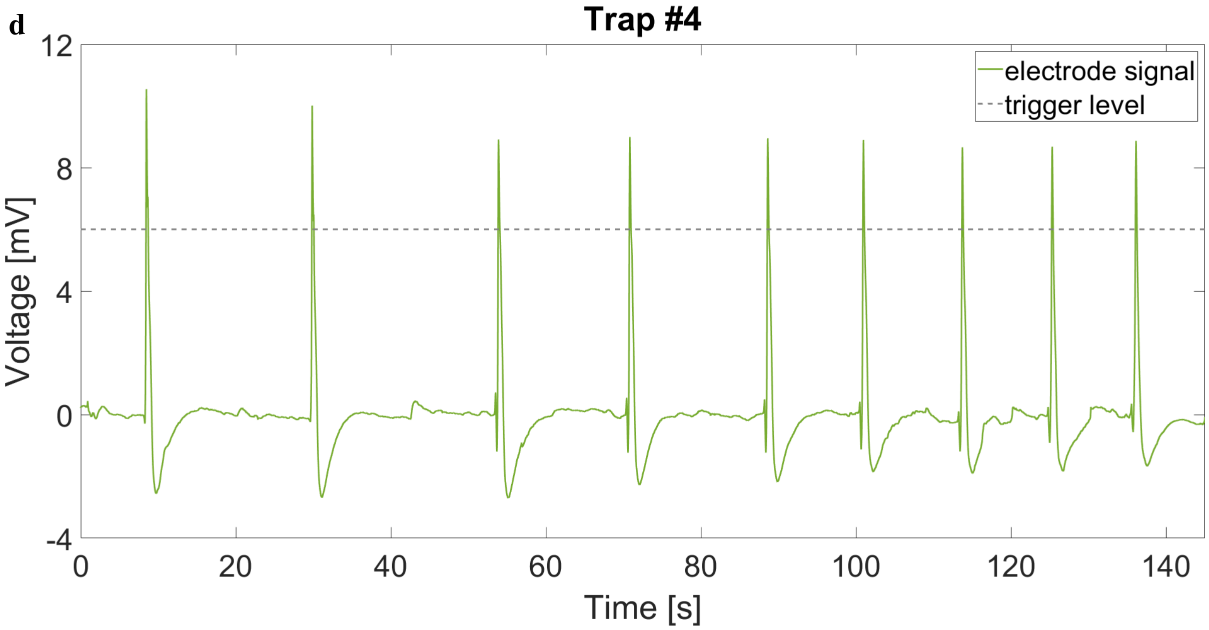


**Extended Data Fig. 5 | Electric time traces showing heat-induced action potentials in four separate experiments with different plant samples.** Corresponds to the data shown in Fig. 5 of the main text. The APs are used as a trigger so that we can perform averaging of the simultaneous magnetic data; the trigger level is indicated by the gray dashed line.

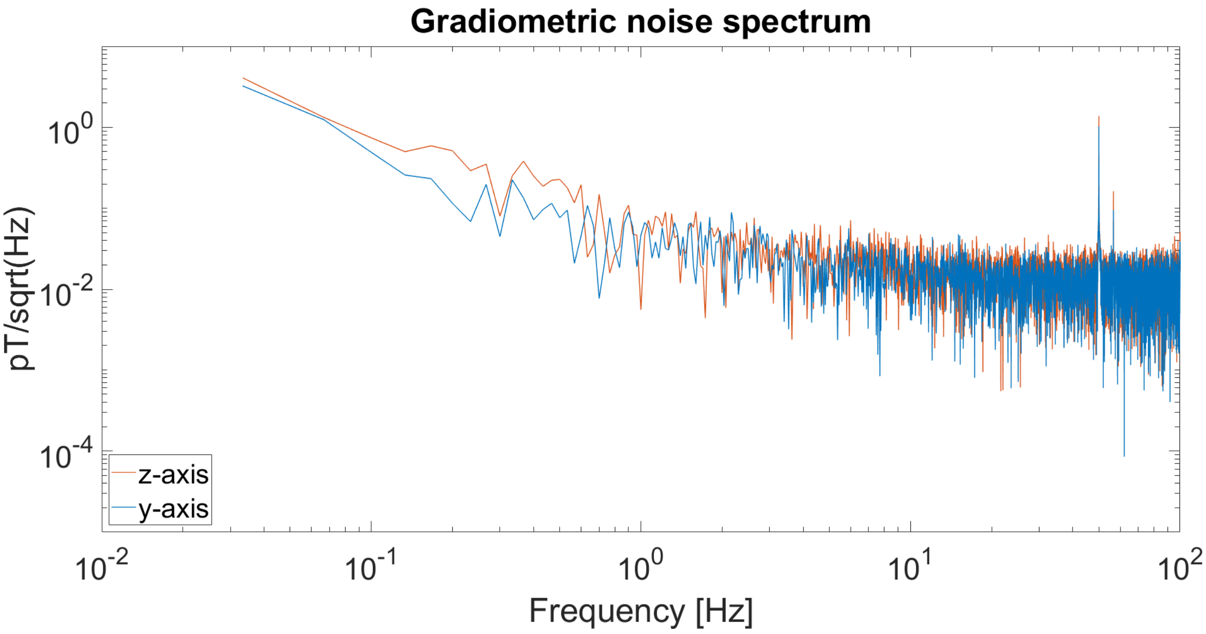


**Extended Data Fig. 6 | Typical noise floor of the gradiometer in the magnetically shielded room.** Obtained by recording a 30-s time trace prior to the start of an experiment.
